# SARS-CoV-2 D614 and G614 spike variants impair neuronal synapses and exhibit differential fusion ability

**DOI:** 10.1101/2020.12.03.409763

**Authors:** Chiung-Ya Chen, Yu-Chi Chou, Yi-Ping Hsueh

## Abstract

Severe acute respiratory syndrome coronavirus 2 (SARS-CoV-2) that causes Coronavirus disease 2019 (COVID-19) exhibits two major variants based on mutations of its spike protein, i.e., the D614 prototype and G614 variant. Although neurological symptoms have been frequently reported in patients, it is still unclear whether SARS-CoV-2 impairs neuronal activity or function. Here, we show that expression of both D614 and G614 spike proteins is sufficient to induce phenotypes of impaired neuronal morphology, including defective dendritic spines and shortened dendritic length. Using spike protein-specific monoclonal antibodies, we found that D614 and G614 spike proteins show differential S1/S2 cleavage and cell fusion efficiency. Our findings provide an explanation for higher transmission of the G614 variant and the neurological manifestations observed in COVID-19 patients.

## Introduction

COVID-19 caused by SARS-CoV-2 infection (Lu et al., 2020; Wu et al., 2020) represents an unprecedented threat to global health and economics. Apart from developing vaccines and rapid screening systems, it is also urgent to understand characteristics of SARS-CoV-2 (SARS2 in short hereafter) that may inform possible therapeutics for the COVID-19 pandemic.

SARS2 binds to the receptors angiotensin-converting enzyme 2 (ACE2) and neuropilin-1 (NRP1) on host cell surfaces via its spike envelope glycoprotein, triggering membrane fusion for viral entry (Cantuti-Castelvetri and al., 2020; Daly and al., 2020; Lan et al., 2020; Li, 2016; Lu et al., 2020; Walls et al., 2020; Yan et al., 2020). In addition to viral entry, SARS2 spike protein also triggers fusion of host cells, giving rise to giant cells harboring multiple nuclei (also known as syncytia) in patients (Ou et al., 2020). Since ACE2 is widely distributed in divergent cell types, including neurons and glial cells (Chen et al., 2020b; Song et al., 2020), like SARS-CoV (SARS in short hereafter) (Guo et al., 2008), SARS2 can infect multiple organs, including the olfactory and central nervous systems (Baig and Sanders, 2020; Bullen et al., 2020; Butowt and Bilinska, 2020; Ellul et al., 2020; Iadecola et al., 2020; Li et al., 2020a; Li et al., 2020c; Natoli et al., 2020; Paniz-Mondolfi et al., 2020; Varatharaj et al., 2020; Zhou et al., 2020a; Zubair et al., 2020). Human brain organoid cultures have further confirmed that SARS2 is able to infect human cortical neurons, evidenced by the presence of respective viral genomes and proteins in neurons (Bullen et al., 2020; Mesci and al., 2020; Song et al., 2020). Studies of animal models also support that both SARS and SARS2 can infect neurons of the rodent central nervous system (CNS) (Natoli et al., 2020; Netland et al., 2008). SARS2 infection results in neurological manifestations and neuropsychiatric illness in both adult and newborn patients, including loss of olfaction and/or taste, dizziness, headache, impaired consciousness, delirium/psychosis, seizures, inflammatory CNS syndromes, and rare cases of ataxia and epilepsy (Chen et al., 2020a; Filatov et al., 2020; Li et al., 2020c; Mao et al., 2020a; Mao et al., 2020b; Paniz-Mondolfi et al., 2020; Paterson et al., 2020; R.W. and al., 2020; Zhou et al., 2020b; Zubair et al., 2020). Neurological symptoms persist in COVID-19 long-haulers, who still have the signs of damage after recovery from infection (Marshall, 2020; Weerahandi and al., 2020), indicating long-term impacts of COVID-19 on the nervous system. Neuronal cell death and reduced protein levels of neuronal markers have been reported in SARS2-infected human brain organoids (Mesci and al., 2020; Song et al., 2020). However, it remains unclear if SARS2 influences neuronal function or activity to cause neurological symptoms. Recent proteomic studies have analyzed protein-protein interactions among SARS2 viral proteins and host proteins (Gordon and al., 2020) and the phosphorylation landscape of SARS2-infected cells (Bouhaddou et al., 2020). These studies indicate that SARS2 viral proteins interact with host proteins, thereby altering cellular status.

Mutations in SARS2 spike protein influence viral infectivity (Li et al., 2020b). In particular, the D614G mutation of SARS2 spike resulted in a global transition of SARS2 from the D614 prototype originally identified from Wuhan to the G614 variant that became dominant since late May 2020 (Korber et al., 2020; Li et al., 2020b). To date, it is unclear why the D614G mutation enhances SARS2 infectivity (Becerra-Flores and Cardozo, 2020; Daniloski et al., 2020; Korber et al., 2020; Zhang et al., 2020). In this report, we used primary cultured neurons and three different cell lines to investigate biochemical and cell biology properties of spike D614 and G614 variants. We unexpectedly found that both D614 and G614 spike proteins alter neuronal morphology. Moreover, these two spike variants have differential properties in terms of S1/S2 proteolysis and cell fusion, which may account for variant infectivity of SARS-CoV-2 variants.

## Results

### Expression of D614 spike alters dendritic spine density and morphology of neurons

We first used cultured neurons as a model to investigate if expression of spike protein is sufficient to alter neuronal morphology, which is highly relevant to neuronal function (Chen et al., 2011; Zoghbi and Bear, 2012). We co-transfected HA-tagged D614 spike proteins and GFP into cultured neurons at 13 days in vitro (DIV) and harvested neurons for immunostaining at DIV 18, representing the stage when cultured neurons are sufficiently mature to form synapses (Chen et al., 2017; Hung et al., 2018). GFP signals were used to outline neuronal morphology, including dendritic spines, the subcellular structures harboring the excitatory synapses of mammalian brain (Harris and Stevens, 1989). We found that spike protein, as revealed by HA immunoreactivity, was widely distributed in cultured neurons, including their dendritic spines (**Fig. 1A**). A quantification analysis indicated that spike protein increased dendritic spine density, but narrowed spine heads and extended spine length (**Fig. 1B, 1C**), both of which are morphological features of spines having functional defects (Hering and Sheng, 2001; Hsueh, 2012). These results indicate that expression of SARS2 spike protein likely impairs the synaptic function of host neurons, potentially giving rise to neurological symptoms in patients.

**Figure 1.**
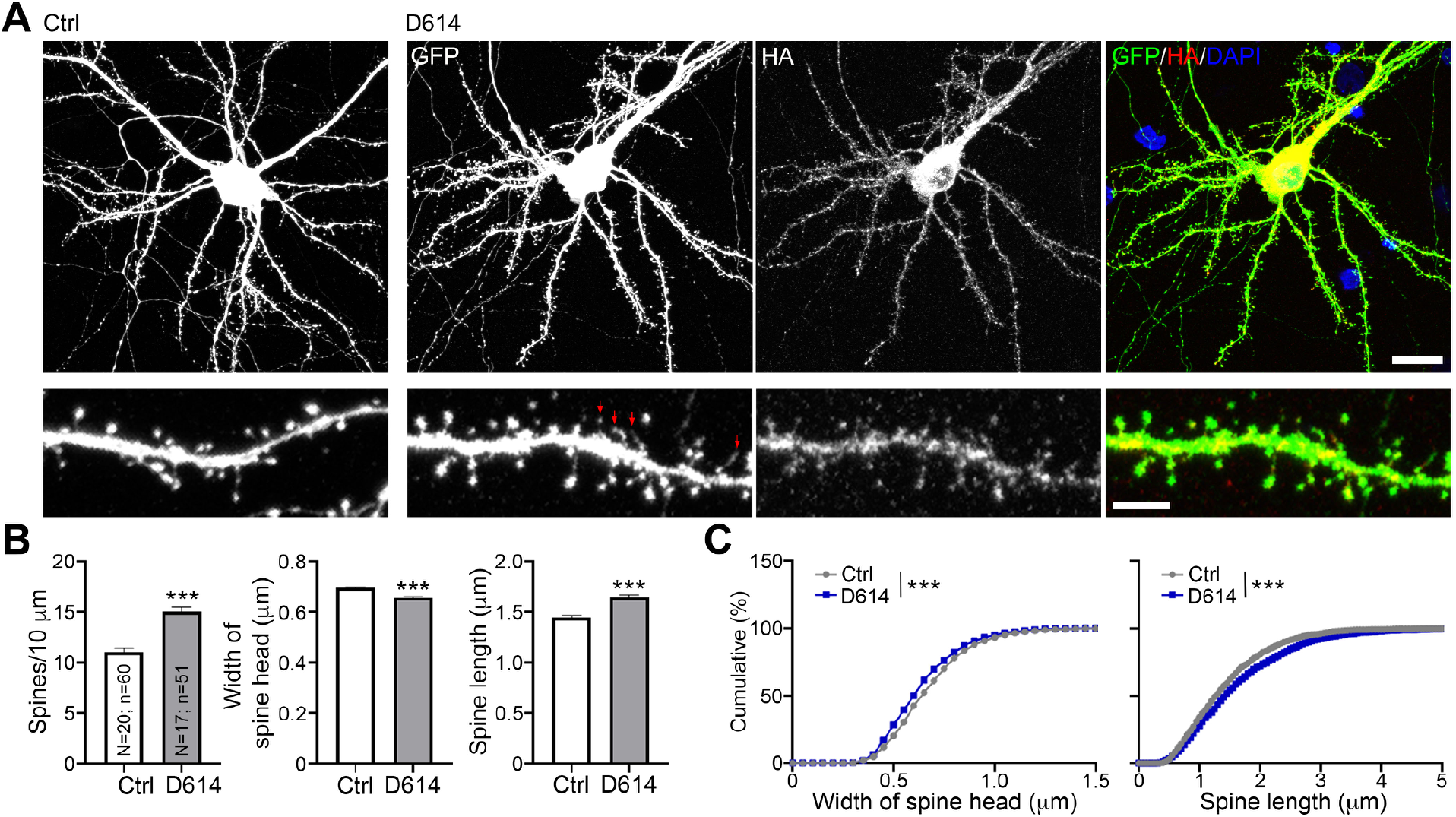
Overexpression of SARS2 spike protein alters the density and morphology of dendritic spines of cultured neurons. (**A**) Mouse cortical neurons were cotransfected with GFP and either vector control or HA-tagged D614 spike protein at DIV13. Five days later, immunostaining was performed to monitor GFP and HA signals. Counterstaining with DAPI was included to label the nuclei of neurons. Red arrows point to some very tiny spine heads. Scale bar: upper, 20 μm; lower, 5 μm. (**B**) Quantification of spine density, length and width. The results represent mean plus SEM. (**C**) Cumulative probability of the width and length of dendritic spines. The experiments were independently repeated three times. Only one set of representative data is shown. The numbers of examined neurons (N) and dendrites (n) are shown in columns. Three clearly identified dendrites of each neuron were subjected to analysis of dendritic spine density and morphology. ***, *P* < 0.001.

### Generation and characterization of SARS2 spike monoclonal antibodies

To further characterize SARS2 spike proteins, we used the receptor binding domain (RBD) of SARS2 spike protein as an immunogen to generate mouse monoclonal antibodies (**Fig. S1A**). Seven specific mouse monoclonal antibodies for the RBD of SARS2 spike protein were generated, only two of which (namely SA07 and SA15) were used in this report. Our RBD monoclonal antibodies recognized their immunogen (i.e. MBP-RBD fusion protein) as well as C-terminal HA-tagged full length (FL) spike proteins expressed in human embryonic kidney HEK239T cells (**Fig. S1B)**. Note that, in addition to the C-terminal HA tagging, we did not introduce any other mutation in our SARS2 spike protein to prevent it from proteolytic process and ER retention. Thus, these two RBD antibodies also detected a 120-kDa band that presumably represents the S1 fragment of SARS2 spike proteins after cleavage. Notably our antibodies did not recognize SARS spike proteins (**Fig. S1B**), evidencing their specificity. On the other hand, HA and SARS-CoV S2 (CoV-S2) antibodies detected FL and S2 fragments (a protein band at ~105 kDa) of SARS2 spike (**Fig. S1B**). Apart from in denatured conditions, our RBD antibodies also worked well in other applications to detect native spike protein, i.e., enzyme-linked immunosorbent assay (ELISA) (**Fig. S1C**), immunoprecipitation (**Fig. S1D**), and immunostaining (**Fig. S1E, S1F**). Thus, our two newly generated RBD antibodies (SA07 and SA15) can serve as powerful tools in multiple assays to enable investigations of the expression, subcellular distribution, and proteolysis of SARS2 spike protein in host cells.

### Differential proteolysis and cell fusion activity of D614 and G614 spike proteins

Sine the S2 fragment generated by proteolytic cleavage of furin is essential for viral entry and cell fusion mediated by SARS-CoV-2 spike proteins, we speculate that the D614G mutation alters proteolytic processing of SARS2 spike protein and, consequently, enabled the change in viral infectivity. To investigate that possibility, we used three different cell lines to express SARS2 spike proteins, namely mouse neuroblastoma Neuro-2A cells, African green monkey kidney fibroblast-like COS-1 cells and HEK293T cells, and used immunoblotting to assess SARS2 D614 and G614 spike variants. We first observed that the efficiency of S1/S2 proteolysis varied among the tested cell lines (**Fig. 2A–2C**). The G614 variant seemed to be expressed higher than D614 prototype in Neuro-2A cells, though this difference was not obvious in COS1 and HEK293T cells (**Fig. 2A–2C**). Since FL spike proteins are synthesized at endoplasmic reticulum (ER) and then transported to Golgi apparatus for furin cleavage (de Haan et al., 2004; Follis et al., 2006; Hasan et al., 2020; Hoffmann et al., 2020), thereby generating the S1 and S2 fragments, the distinct amounts of S1 and S2 fragments in different cell lines indicates that these cells exhibit differential ER-Golgi trafficking or furin activities. Thus, our data imply that different cell types show differential abilities to process SARS2 spike proteins.

**Figure 2.**
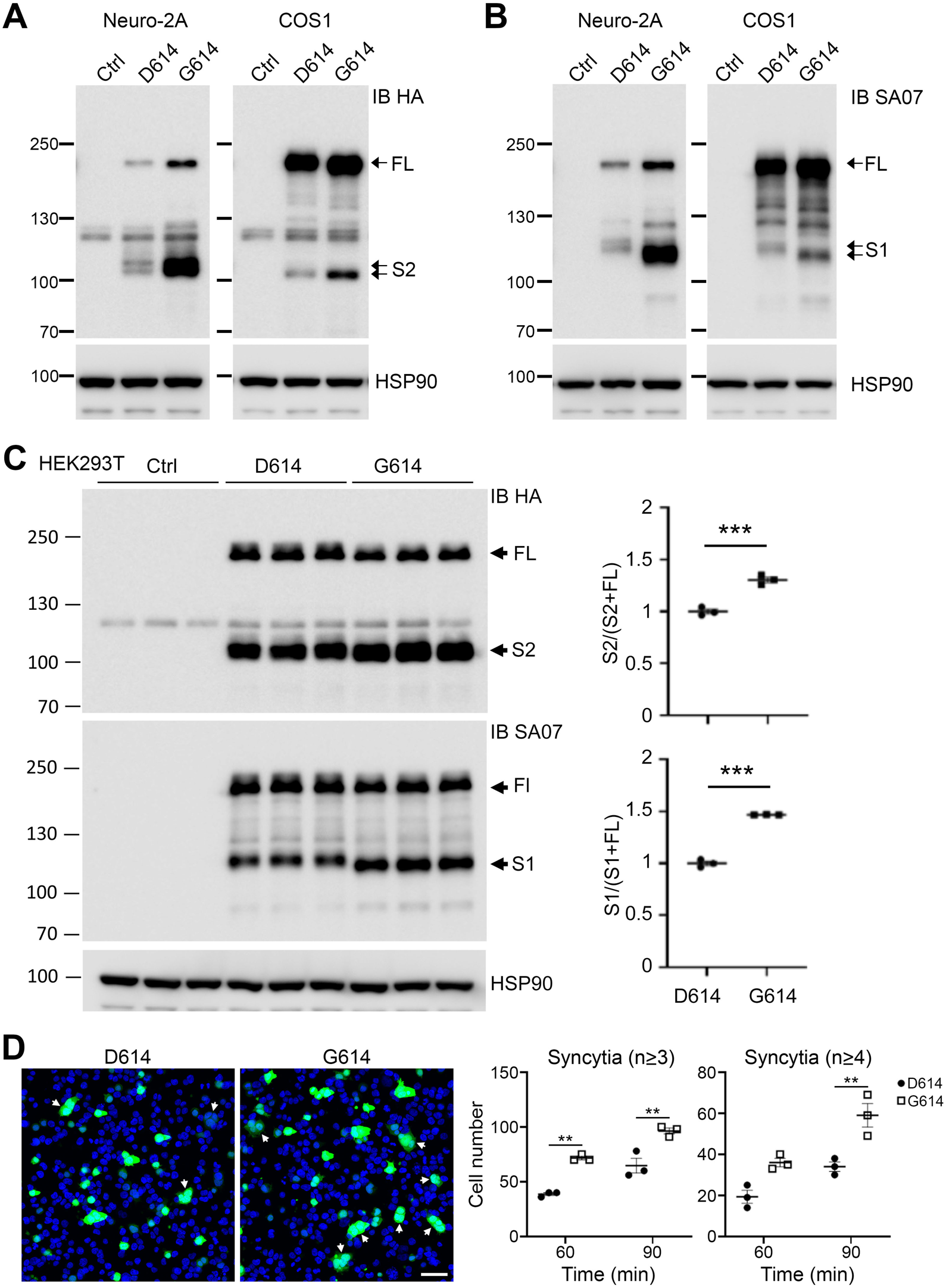
D614 and G614 spike proteins show distinct patterns of proteolysis and cell fusion ability. (**A)-(C)** Neuro-2A, COS1 and HEK293T cells were transfected with vector control and HA-tagged spike proteins (D614 and G614) and analyzed using immunoblotting with HA and SA07 antibodies as indicated. FL, full length spike protein; S1 and S2, the cleaved products of spike protein; HSP90, internal loading control. Note several potential glycosylation variants were present. HA antibody exhibited crossreactivity with protein bands of ~125 kDa in the control group. However, SA07 antibody did not present any signal in the control group. In (**C**), cell lysates of each group were collected from three different culture wells. The results of quantification are also summarized. (**D**) Cell-cell fusion mediated by spike protein and hACE2. HEK293T cells were transiently expressed with GFP and either D614 or G614 spike protein. The cells were then detached using EDTA, and co-cultured with 293T/hACE2 cells for 60 or 90 min at 37°C. Enlarged GFP-positive cells possessing three or more nuclei were counted (white arrows). “Syncytia (n>3)” represents GFP-positive cells containing at least three nuclei. “Syncytia (n>4)” represents GFP-positive cells with four or more nuclei. Experiments were independently repeated three times. The results represent mean plus SEM. **, *P* < 0.01; ***, *P* < 0.001. Scale bar, 50 μm.

Intriguingly, we found that proteolysis of the G614 variant was greater than that of the D614 prototype, as S2 fragments of G614 spike proteins were relatively more abundant than those of D614 spike proteins in all the tested cell lines (**Fig. 2A–2C**). In addition, we observed that some G614 variants (including FL, glycosylation variants and S1 fragments) migrated slightly faster by SDS-PAGE relative to the D614 prototype (**Fig. 2A–2C**). Apart from being glycosylated (Watanabe et al., 2020), spike proteins are posttranslationally modified by phosphorylation (Bouhaddou et al., 2020), which likely contributes to the differing SDS-PAGE mobility of the D614 and G614 variants.

Since the cleaved S2 fragment mediates membrane fusion, the higher levels of S2 fragments derived from G614 spike protein support our speculation that G614 spike protein may present better fusion ability relative to D614 spike. To confirm that point, we investigated the syncytium formation ability of D614 and G614 spike proteins. SARS2 spike protein and human ACE2 were separately expressed in two different HEK293T cell cultures. GFP was cotransfected with spike protein to monitor cell fusion. Sixty and ninety minutes after adding spike/GFP-expressing cells to human ACE2-expressing cells, the cells were fixed to monitor cell fusion. Enlarged GFP cells possessing three or more nuclei were identified as syncytia (**Fig. 2D,** arrows). For both 60- and 90-min co-culture, numbers of syncytia were higher for G614-relative to D614-expressing cells (**Fig. 2D**, middle). Numbers of syncytia having ≥4 nuclei were also noticeably higher in the group of G614-expressing cells after 90-min co-culture (**Fig. 2D**, right). Both spike protein and ACE2 were required for cell fusion because neither spike protein alone nor ACE2 alone induced cell fusion (data not shown). These results support that the G614 variant is more effective at cell fusion than the D614 prototype. Moreover, they imply that the SARS2 G614 variant endows more effective viral entry and triggers greater cell fusion of infected cells with neighboring non-infected cells. The fact that G614-mediated transmission is more effective may account for the global transition from the D614 to G614 variant (Korber et al., 2020).

### Differential S1 and S2 epitope distribution of spike proteins

Next, we performed fluorescence immunostaining to further characterize spike protein expression in host cells. Spike protein targets to the dendritic spines of neurons (**Fig. 1A**), which are F-actin-enriched structures. Therefore, we investigated if spike proteins also colocalize with F-actin in HEK293T and Neuro-2A cells. Z-series confocal imaging clearly indicated strong HA or MYC immunoreactivities of both D614 and G614 variants in F-actin-related structures, i.e. filopodia, lamellipodia, cell edges and stress fibers (**Fig. 3A, 3B**). However, SA07 antibody revealed a mainly intracellular pattern for both D614 and G614 spike proteins in HEK293T and Neuro-2A cells (**Fig. 3C, 3D**). Enrichment of spike protein variants in F-actin-positive structures was not universal for other membrane proteins, as Toll-like receptor 3 (TLR3) was not concentrated at filopodia or lamellipodia in HEK293T cells (**Fig. S2**). As spike protein contains an ER retention signal, next we examined if the intracellular SA07-staining pattern was indeed associated with ER. We found that SA07 immunoreactivities were considerably colocalized with the ER marker calreticulin (Calr) in both D614- and G614-expressing HEK293T cells, though SA07 immunoreactivity extended further to cell margins, i.e., a region lacking calreticulin (**Fig. 4A, 4B**, middle panel, red only). However, we found that neither MYC nor CoV-S2 antibody presented obvious colocalization with calreticulin (**Fig. 4A, 4B**, upper and lower panel). Instead, these antibodies clearly outlined cell margins and also labeled intracellular aggregates (**Fig. 4A, 4B**). That outcome was not specific to SARS2 spike protein because the same pattern was observed for SARS spike protein (**Fig. 4C**). It was also not specific for HEK293T cells because a similar distribution was apparent for Neuro-2A cells (**Fig. 4D**). Since HA, MYC and CoV-S2 antibody all detect FL and S2 fragments of spike protein and SA07 recognizes the FL and S1 fragments, these observations indicate that the subcellular distribution of the S1 and S2 fragments of spike protein differ. The S1 epitope was highly enriched at ER and the S2 epitope tended to target to the cell surface and filopodia. The occurrence of S2 fragments alone in filopodia may further promote the cell fusion efficiency of infected cells with neighboring cells (**Fig. 3**). In the context of COVID-19 patients, this S2 distribution would also be expected to enhance syncytia formation to impair cellular function.

**Figure 3.**
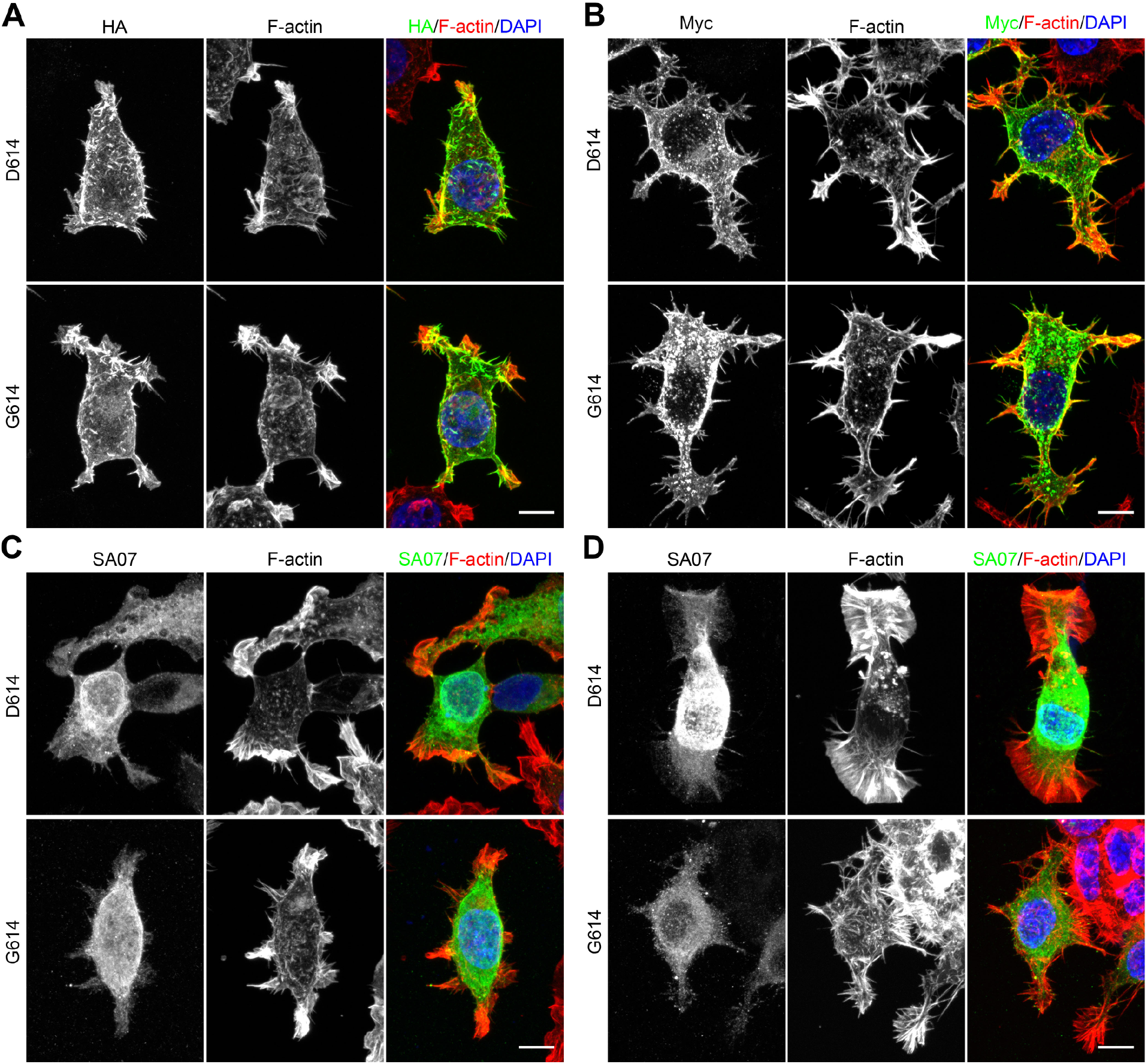
The S1 and S2 fragments of SARS2 spike protein exhibit differential distributions in cells. (**A, C**) HA-tagged and (**B, D**) MYC-tagged D614 and G614 constructs were transfected into (**A, C**) HEK293T cells or (**B, D**) Neuro-2A cells for immunostaining using HA, MYC and SA07 antibodies. F-actin was labeled by Alexa-fluor-546-conjugated phalloidin. DAPI was used to label the nuclei. Scale bars: 10 μm.

**Figure 4.**
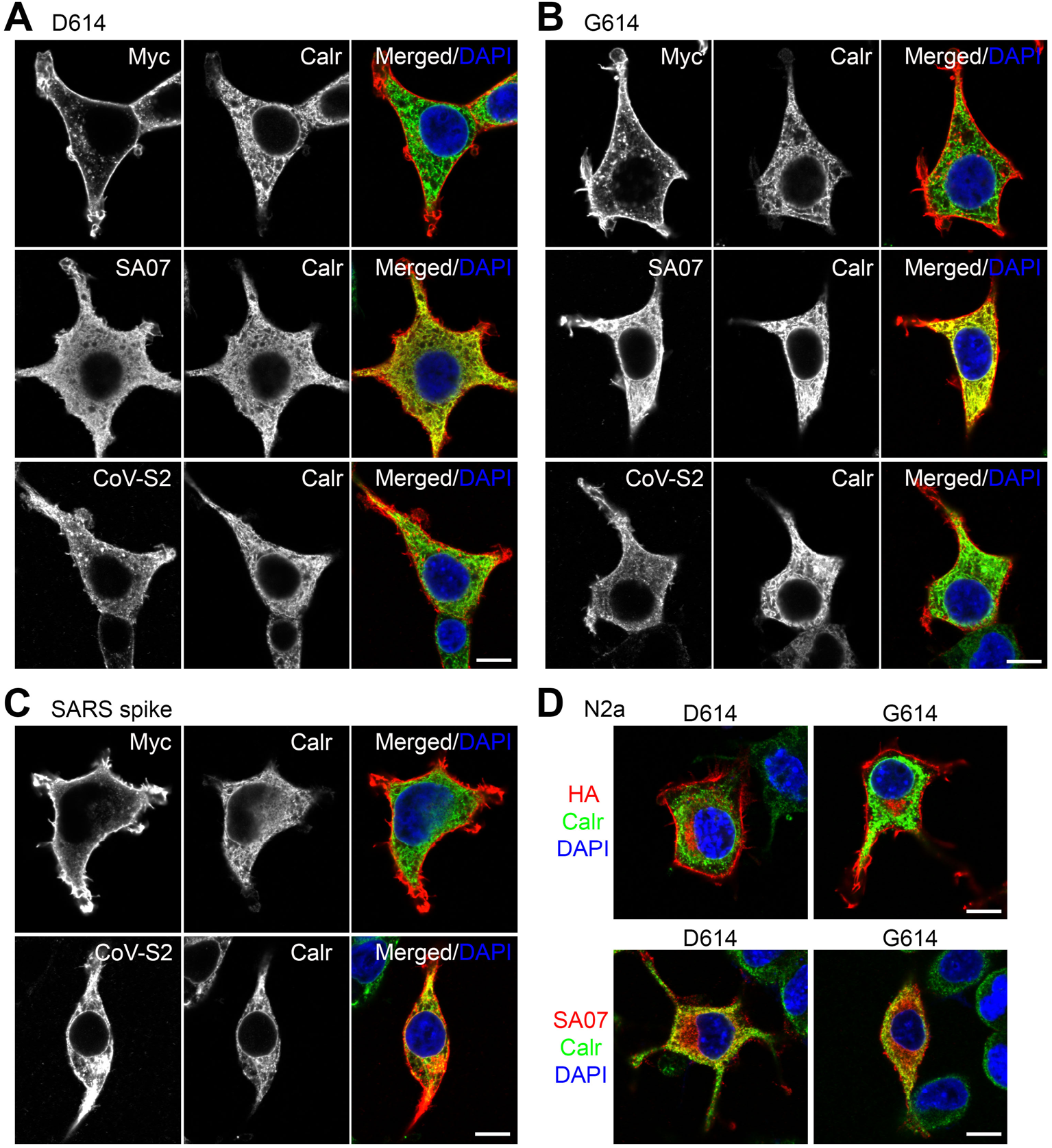
RBD and S2 immunoreactivities are differentially distributed across the plasma membrane and ER. (**A)-(C)** HEK293T and (**D**) Neuro-2A cells were transfected with various plasmids as indicated. Dual immunostaining using the ER marker calreticulin (Calr) and MYC, SA07 and CoV-S2 antibodies was performed and analyzed by means of confocal microscopy. The individual and merged images are shown. Counterstaining using DAPI to label the nuclei was also performed. Scale bars: 10 μm.

### Both D614 and G614 spike variants alter neuronal morphology

As the S2 fragments of both the D614 and G614 variants targeted to filopodia, we investigated if the G614 variant also results in dendritic spine defects of cultured neurons. We co-transfected GFP with vector control, D614 or G614 into cultured hippocampal neurons at DIV 13. Similar to observations for the D614 prototype, the G614 variant was also widely present in neurons, including in dendritic spines (**Fig. 5A**). Moreover, both the G614 and D614 variants exerted comparable effects in terms of reducing dendritic spine density, narrowing spine heads and extending spine length (**Fig. 5B**). Thus, G614 spike proteins exhibit equivalent ability to the D614 prototype in impairing synaptic activity and morphology. Since SARS2 infects both adult and newborn patients, we also investigated the effect of spike variants on developing neurons (**Fig. 5C, 5D**). Cultured neurons were co-transfected with HA-tagged spike proteins and GFP at DIV 3 and harvested for immunostaining at DIV 6. We found that expression of both SARS2 spike variants shortened dendritic lengths of developing neurons (**Fig. 5C, 5D**). Thus, both D614 and G614 spike proteins have the ability to impair neuronal morphology.

**Figure 5.**
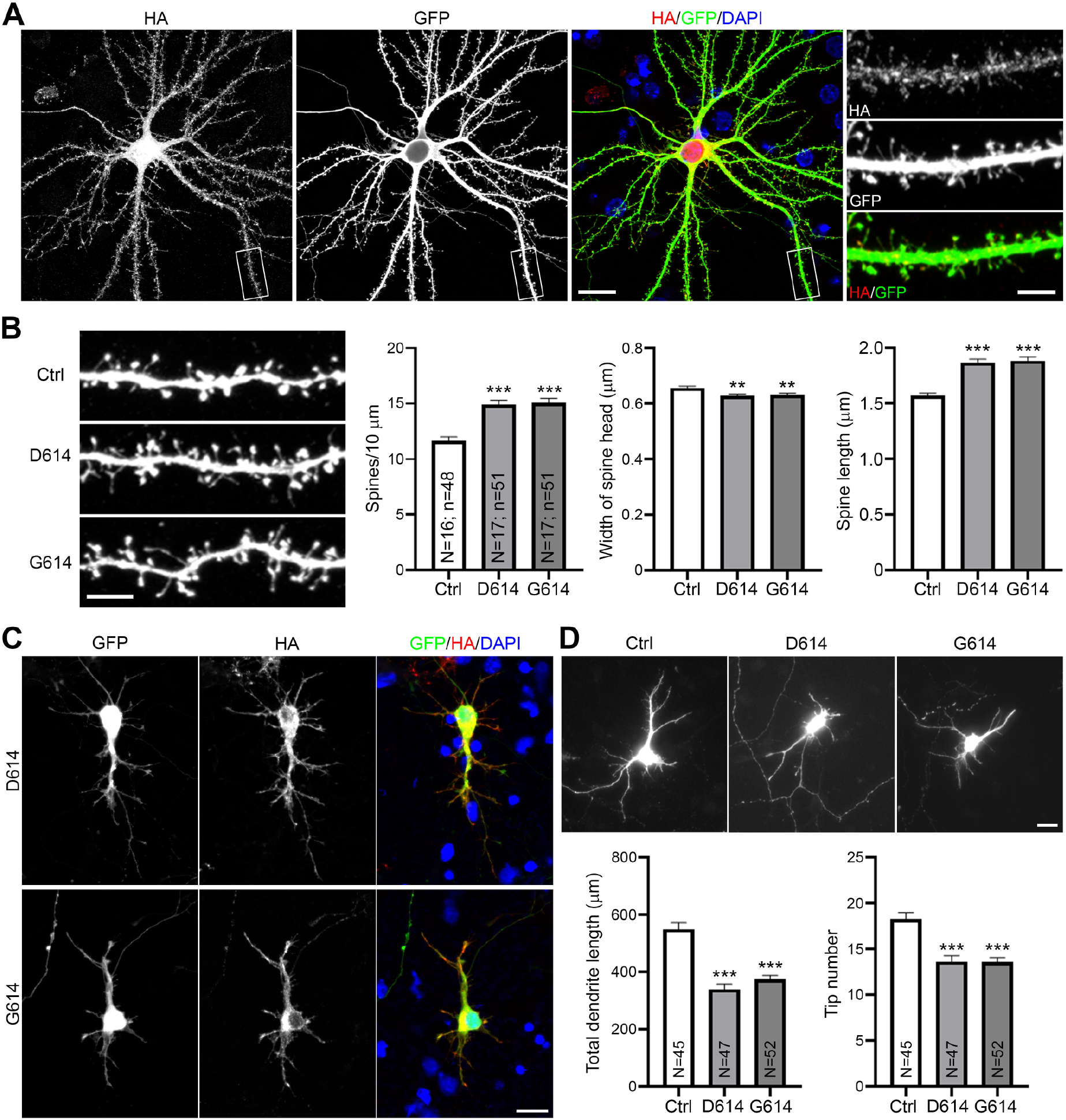
Both D614 and G614 spike proteins induce dendritic spine defects and impair dendritic growth in cultured neurons. (**A**) Representative images of a G614-expressing neuron at DIV 18. The experiment was conducted as described in the legend of Figure 1. (**B**) Quantification of (**A**). (**C**) Representative images of D614 and G614 expressing neuron at DIV6. Neurons were transfected with indicated plasmids and GFP at DIV 3. Neuron morphology was determined by GFP signals. (**D**) Quantification of (**C**). Total dendrite length and the number of dendritic tips were determined to indicate dendritic growth of immature neurons. For (**B**) and (**D**), the experiments were independently repeated three times. Only one set of representative data is shown. The numbers of examined neurons (N) and dendrites (n) are shown in columns. The results represent mean plus SEM. **, *P* < 0.01, ***, *P* < 0.001. Scale bar: (**A**) full image, 20 μm; dendrite segment, 5 μm; (**B**) 5 μm; (**C-D**) 20 μm.

## Discussion

In this report, we show that expression of SARS2 spike protein is sufficient to impair neuronal morphology. Both the D614 and G614 spike variants cause defects in the dendritic spines of mature neurons and reduce the dendritic length of developing neurons. Since neuronal morphology is highly relevant to neuronal function, the morphological defects caused by SARS-CoV-2 spike protein imply that infected neurons function abnormally, which is likely relevant to the neurological and neuropsychiatric symptoms presented by COVID-19 patients (Chen et al., 2020a; Ellul et al., 2020; Filatov et al., 2020; Li et al., 2020c; Mao et al., 2020b; R.W. and al., 2020; Varatharaj et al., 2020; Zhou et al., 2020b). Nevertheless, to confirm altered neuronal activity upon SARS2 infection, it is imperative to perform electrophysiological recordings in the future. Importantly, it is unclear how spike proteins control neuronal morphology. Since spike protein distribution overlaps with that of F-actin and given that F-actin cytoskeletons are critical regulators of neuronal morphology (Dillon and Goda, 2005; dos Remedios et al., 2003), spike proteins may control F-actin dynamics or rearrangement via an unknown mechanism to influence neuronal morphology. It will be intriguing to investigate the mechanism by which spike proteins might target to F-actin and regulate F-actin dynamics.

It has been questioned if SARS2 can infect neurons by binding to ACE2 because ACE2 expression levels are low in the nervous system (Cai et al., 2020; Cooper et al., 2020). We postulate three possible mechanisms by which SARS2 can infect neurons. First, though low, *ACE2* transcripts are still detectable in neurons (Chen et al., 2020b). Low-level ACE2 expression may be sufficient for infection when viral titers are high. Secondly, infection may be mediated by another spike receptor, NRP1, which is highly expressed in neurons (He and Tessier-Lavigne, 1997; Kawakami et al., 1996; Kolodkin et al., 1997). Thirdly, membrane fusion may be mediated by digested S2 fragments (Cai et al., 2020), since such fragments are present on virion surfaces (Cai et al., 2020) and SARS2 spike protein-expressing cells (presumably virus-infected cells in patients). Consequently, neurons may be infected by locally high virion dosages released from proximal infected tissues or upon fusing with neighboring infected cells. Certainly, these three possibilities are not mutually exclusive and further investigations are required to elucidate the detailed infection pathways of SARS2 in vivo.

Using a series of biochemical and cell biological studies, we report three differences between D614 and G614 spike proteins. First, our results echo a previous cryo-EM study that spike proteins are readily processed into S1 and S2 fragments in host cells (Cai et al., 2020). S1/S2 cleavage of G614 spike is more pronounced compared to that of the D614 prototype. Second, perhaps due to having higher amounts of S2 fragments, G614 spike protein-expressing cells exhibited more efficient cell fusion than D614-expressing cells. Third, G614 and D614 spike proteins exhibit slightly different mobilities on SDS-PAGE gels, indicating that the D614G mutation may influence posttranslational modifications to modulate spike protein activity. In terms of proteolytic processing of spike variants, the conclusions of several studies are conflicting (Daniloski et al., 2020; Hu and al., 2020; Yurkovetskiy and al., 2020). Using pseudotyped virus bearing SARS2 spike protein, some researchers did not observe distinct proteolytic cleavage between D614 and G614 variants. However, when producing pseudotyped virus, the furin cleavage site and ER-retention signal of spike protein tends to be mutated or removed in order to obtain high concentration virion titers (Ou et al., 2020; Yurkovetskiy and al., 2020). Some structural studies have also used the same strategy to generate high yields of homogenous spike proteins (Yurkovetskiy and al., 2020). Consequently, this kind of modification may alter spike protein properties, contributing to discrepancies among results.

Although infection mechanisms in virus-infected cells are expected to be more complex than for expression of spike protein alone and some key conclusions should be further validated in SARS2-infected cells, our study sheds light on the properties of SARS2 spike protein in host cells. Our findings also provide possible explanations for why the D614G mutation enhances infectivity and how SARS2 infection impairs neuronal function.

## Methods and Materials

### Animals

WT C57BL/6 and BALB/c mice were purchased from the National Laboratory Animal Center, Taipei, Taiwan. Animals were housed in the animal facility of the Institute of Molecular Biology, Academia Sinica, with a 12-h light/12-h dark cycle and controlled temperature and humidity. All animal experiments were performed with the approval of the Academia Sinica Institutional Animal Care and Utilization Committee and in strict accordance with its guidelines and those of the Council of Agriculture Guidebook for the Care and Use of Laboratory Animals. For primary neuronal cultures, E16-17 C57BL/6 mouse embryos of both sexes were used. For SARS-CoV-2 spike RBD antibody generation, only male BALB/c mice were used.

### Constructs

Plasmid *UC57-2019-nCoV-S* (Human) was kindly provided by Dr. Yu-Chan Chao, Institute of Molecular Biology, Academia Sinica. The receptor binding domain (RBD) of SARS-CoV-2 spike (amino acids (aa) 319-541) was amplified by using the primer pairs: RBD Fw-5’ GGGATCCAGGGTGCAGCCAACC 3’ and RBD Rv-5’ GCTGGTCGACTTAGAAGTTCACGCACTTG 3’ and subcloned into pMAL-c2x and pGEX-4T1 vectors at BamHI and SalI sites. Full-length SARS-CoV-2 spike protein (aa 1-1273) was amplified by using the primer pairs: CoV2 Fw-5’ GAAGATCTATGTTCGTCTTCCTG 3’ and CoV2 Rv-5’ GCTGGTCGACGGTGTAATGCAGCTTC 3’ and subcloned into pGw1-cHA and pGw1-cMyc vectors at BglII and SalI sites. Site-directed mutagenesis was performed with the primer pairs: D614G sense-5’ GTGCTGTATCAGGGCGTGAATTGTACC 3’ and D614G antisense-5’ GGTACAATTCACGCCCTGATACAGCAC 3’ to generate the pGw1-SARS2-spike (G614)-HA and pGw1-SARS2-spike (G614)-Myc constructs. pCMV-SARS-spike (codon optimized) was purchased from Sino Biological (VG40150-G-N). Full-length SARS-CoV spike (aa 1-1255) was amplified by using the primer pairs: CoV Fw-5’ GAAGATCTATGTTCATCTTCCTG 3’ and CoV Rv-5’ GCTGGTCGACGGTGTAGTGCAGTTTC 3’ and subcloned into pGw1-cHA and pGw1-cMyc vectors at BglII and SalI sites. Full-length mouse TLR3 (aa 1-905) was amplified by using the primer pairs: Tlr3 Fw-5’ GGGATCCATGAAAGGGTGTTCC 3’ and Tlr3 Rv-5’ GCTGGTCGACATGTGCTGAATTCCG 3’ and subcloned into pGw1-cHA vector at BglII and SalI sites.

### Recombinant protein purification and antibodies

BL21(DE3) *Escherichia coli* bacteria carrying pMAL-c2x-RBD or pGEX-4T1-RBD were grown in LB media with Ampicillin. After OD600 reached 0.5, IPTG was added into the culture to induce MBP-RBD or GST-RBD protein expression. Bacteria were harvested after 3-h induction, and then lysed by sonication. The Triton X-100-extracted soluble fraction was collected, and then the MBP-RBD and GST-RBD proteins were purified by amylose resin and GST beads, respectively. Purified MBP-RBD proteins were used to immunize BALB/c mice. Anti-SARS2-RBD mouse monoclonal antibodies (SA07 and SA15) were generated as described previously (Chen et al., 2011). The HSP90 rabbit polyclonal antibody was a gift from Dr. Chung Wang (Liou and Wang, 2005). The following commercialized antibodies were also used in this study: HA (rabbit, C29F4, Cell Signaling), HA (mouse, 16B12, abcam), Myc (mouse, 9B11, Cell Signaling), GFP (chicken, Abcam), calretinculin (rabbit, ThermoFisher), SARS-CoV/SARS-CoV-2 Spike S2 (mouse, 1A9, GeneTex), anti-mouse-horseradish peroxidase (HRP) (Jackson Lab), anti-rabbit-HRP (GE Healthcare), and anti-mouse/rabbit/chicken-Alexa 488/555 (Invitrogen).

### Cell culture, transfection, and immunoblotting

HEK293T, Neuro-2A, and COS1 cells were cultured in DMEM with 10% FBS plus penicillin and streptomycin at 37 ^o^C and 5% CO_2_. Transfection was performed using Lipofectamine 2000 (Invitrogen) according to the manufacturer’s instructions. To generate the 293T/hACE2 stably-expressed cell line, the human ACE2 gene was amplified from the MGC human cDNA library and sub-cloned into NheI and EcoRI sites of a lentivector provided by the RNAi core facility in Academia Sinica, Taiwan. HEK293T cells were transduced with VSV-G pseudotyped lentivirus carrying the human ACE2 gene and continuously selected with 2.5 mg/mL puromycin. Expression of human ACE2 protein was confirmed by immunoblotting using ACE2 antibody. Mouse cortical neurons were cultured in Neurobasal medium/DMEM (1:1) with B27 supplement and transfected using calcium phosphate precipitation methods as described previously (Hung et al., 2018). Cotransfected GFP was used to outline neuronal morphology. Immunoprecipitation and immunoblotting were performed as described previously (Chen et al., 2017).

### ELISA

A Nunc 96-well microplate (Thermo Scientific) was coated with purified GST-RBD proteins following 2-fold serial dilution starting from 200 ng/ml and incubated at 4 ^o^C overnight. After blocking with 3% BSA in PBS, 1 ug/ml of anti-RBD antibodies (SA07 and SA15) and GST antibody in blocking buffer were added to the wells. After incubation for 1 h, each well was washed five times with 0.1% Tween-20 in phosphate buffered saline (PBS) before applying HRP-conjugated secondary antibody. The plate was incubated at room temperature for another one hour and washed as described above. Substrate solution (0.1 mg/ml 2,2’-azino-bis (3-ethylbenz thiazoline-6-sulphonic acid) (ABTS) and 0.03% H_2_O_2_ in 0.1 M Citric acid pH 4.0) was added into the wells for color development. After 10 to 20 min, the signal of each well was detected using an ELISA reader at OD_405_. For all the solutions used here, the volume applied into each well was 0.1 ml.

### Cell fusion

Spike (D614)-HA or Spike (G614)-HA was co-transfected with GFP into HEK293T cells. After 16-18 h post-transfection, the cells were detached by using 1 mM EDTA/PBS, and then added into 293T/hACE2 cell culture. Cells were fixed after 60 and 90 min co-culture. DAPI was used to counterstain the nuclei. Ten randomly selected non-overlapping areas of each sample were captured using a confocal microscope (LSM 700, Zeiss) equipped with a 20×/NA 0.8 (Plan-Apochromat) objective lens and Zen acquisition and analysis software (Zeiss). Quantitation of fused cells was performed using ImageJ software. An enlarged GFP-positive cell with three or more nuclei was identified as a syncytium.

### Immunostaining

After 24-48 h post-transfection, cells were fixed with 4% paraformaldehyde in PBS at room temperature for 10 min. Neurons were fixed at DIV6 and DIV18. Immunostaining procedures were performed as described previously (Chen et al., 2011). Immunofluorescence images of cultured neurons at DIV6 were visualized with a fluorescence microscope (Axioimage M2; Zeiss) equipped with a 20×/NA 0.8 (Plan-Apochromat) objective lens and acquired using a cooled charge-coupled device camera (Rolera EM-C2; QImaging) with Zen software (Zeiss). For cell lines and DIV18 neurons, images were captured with a confocal microscope (LSM 700, Zeiss) equipped with a 63×/NA 1.4 (Plan-Apochromat) objective lens and Zen acquisition and analysis software (Zeiss). The images were processed using Photoshop (Adobe) with minimal adjustment of brightness or contrast applied to the entire images. Quantitation of neuronal morphology was performed using ImageJ software.

### Neuronal morphometry

For dendrite analysis, total dendrite length, including primary dendrites and all dendritic branches, was measured and the number of dendritic tips was counted. For spine analysis, secondary dendrites of DIV18 neurons were chosen to assess spine density, the width of spine heads, and spine length. All experiments were repeated at least three times. For each repeat, at least 20-30 neurons were randomly picked from each group for analysis. The data shown in this manuscript are the representative results of three independent experiments.

### Statistical analysis

Statistical analysis were performed using GraphPad Prism 8 software. Experiments were performed blind by relabeling the samples with the assistance of other laboratory members. For two-group experiments, an unpaired t-test was used. For experiments with more than two groups, one-way ANOVA with *post hoc* Bonferroni correction was applied. For Figure 2C, two-way ANOVA with *post hoc* Bonferroni correction was used. Data are presented as the mean ± SEM.

## Acknowledgments

We thank Dr. Yun-Fen Hung and the COVID-19 Team of Academia Sinica for excellent technical assistance. Dr. John O’Brien conducted English editing, and members of Dr. Yi-Ping Hsueh’s laboratory relabeled samples for blind experiments. This work was supported by the Institute of Molecular Biology, Academia Sinica.

## Author contributions

C.-Y.C., conceptualization, investigation and writing; Y.-P.H., conceptualization, writing, funding acquisition, supervision and project administration. Competing interests:

## Materials and correspondence

All unique/stable reagents generated in this study are available from Y.-P.H. upon completion of a Materials Transfer Agreement of Academia Sinica.

## List of Supplementary Materials

**Supplementary Figures**

**Figure 1.** Characterization of monoclonal antibodies recognizing RBD of SASR2 spike protein.

**Figure S1.** TLR3 is not enriched in F-actin-positive structures.

## Supplementary Figures and Figure Legends

**Figure S1.**
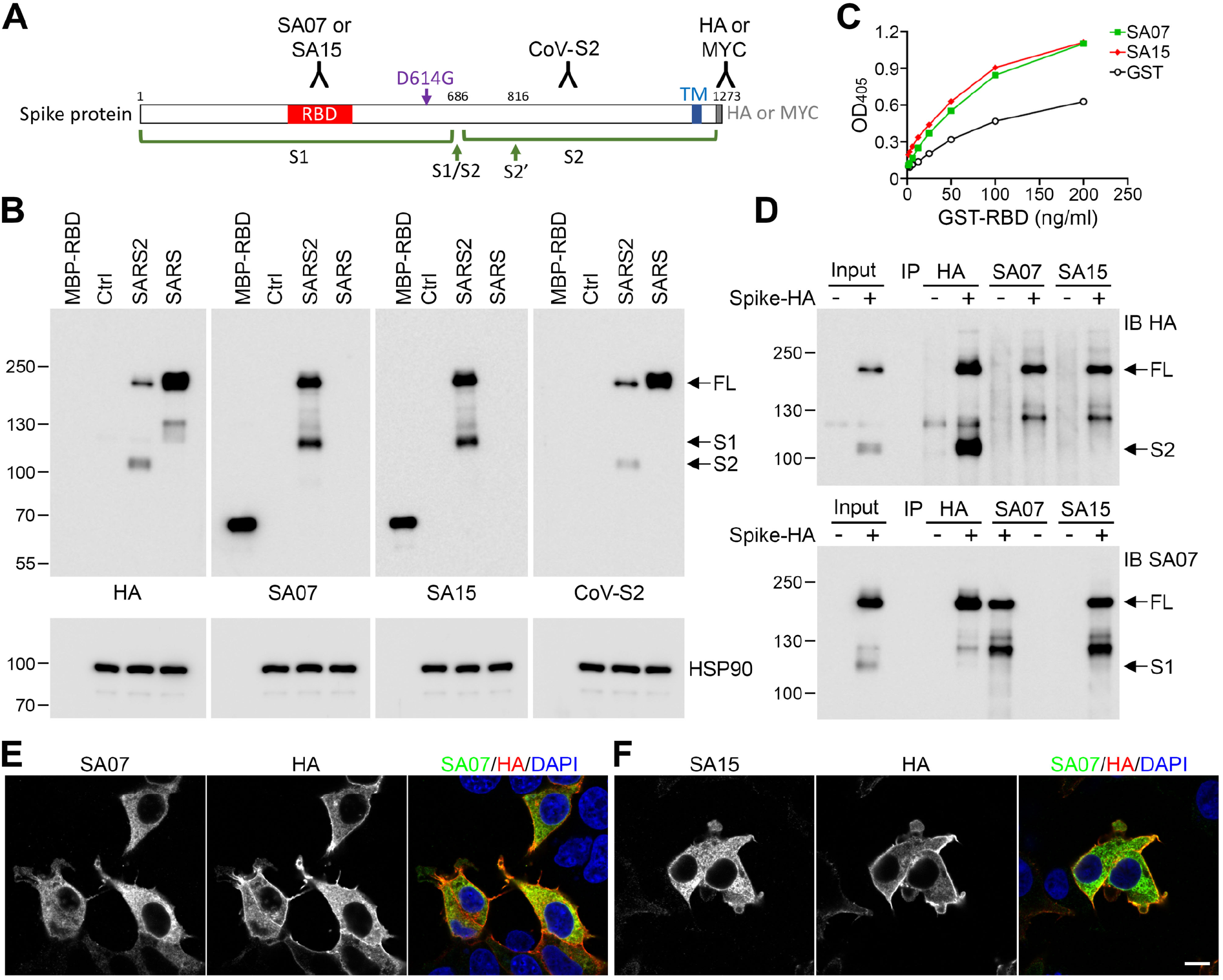
Characterization of monoclonal antibodies recognizing RBD of SASR2 spike protein. (**A**) Schematic domain structure of SARS2 spike protein. Spike protein can be cleaved into S1 and S2 fragments by furin. The D614 residue is located within the S1 fragment. The location of RBD antigens are indicated. Either HA or MYC tagging at the C-terminal of SARS2 spike protein was used in this study. The SARS S2 antibody that is labeled as CoV-S2 can recognize the S2 region of both SARS and SARS2. Thus, the RBD, HA, MYC and CoV-S2 antibodies can be used to label full length, S1 and S2 fragments of SARS2 spike protein. TM, transmembrane domain. (**B**) Immunoblotting analysis reveals the specificity of RBD monoclonal antibodies SA07 and SA15. Purified MBP-RBD fusion protein and HEK293T lysates expressing control vector, SARS2 and SARS spike proteins were included for testing. The results of HA and CoV-S2 antibodies confirmed the expression of SARS and SARS2 spike proteins. HSP90 antibody was used as an internal loading control. (**C**) ELISA using a two-fold series dilution of GST-RBD and SA07, SA15 and GST antibodies. (**D**) Immunoprecipitation of SARS2 spike protein by the RBD antibodies. HEK293T cells were transfected with SARS2 spike protein or control vector. Immunoprecipitation (IP) was performed using SA07, SA15 and HA antibodies. The precipitates were immunoblotted (IB) with HA and SA07 antibodies as indicated. (**E**)-(**F**) Immunostaining using HA tag antibody and SA07 (**E**) and SA15 (**F**) antibodies. Counterstaining using DAPI was performed to label the nuclei. Scale bar: 10 μm.

**Supplementary Fig S2.**
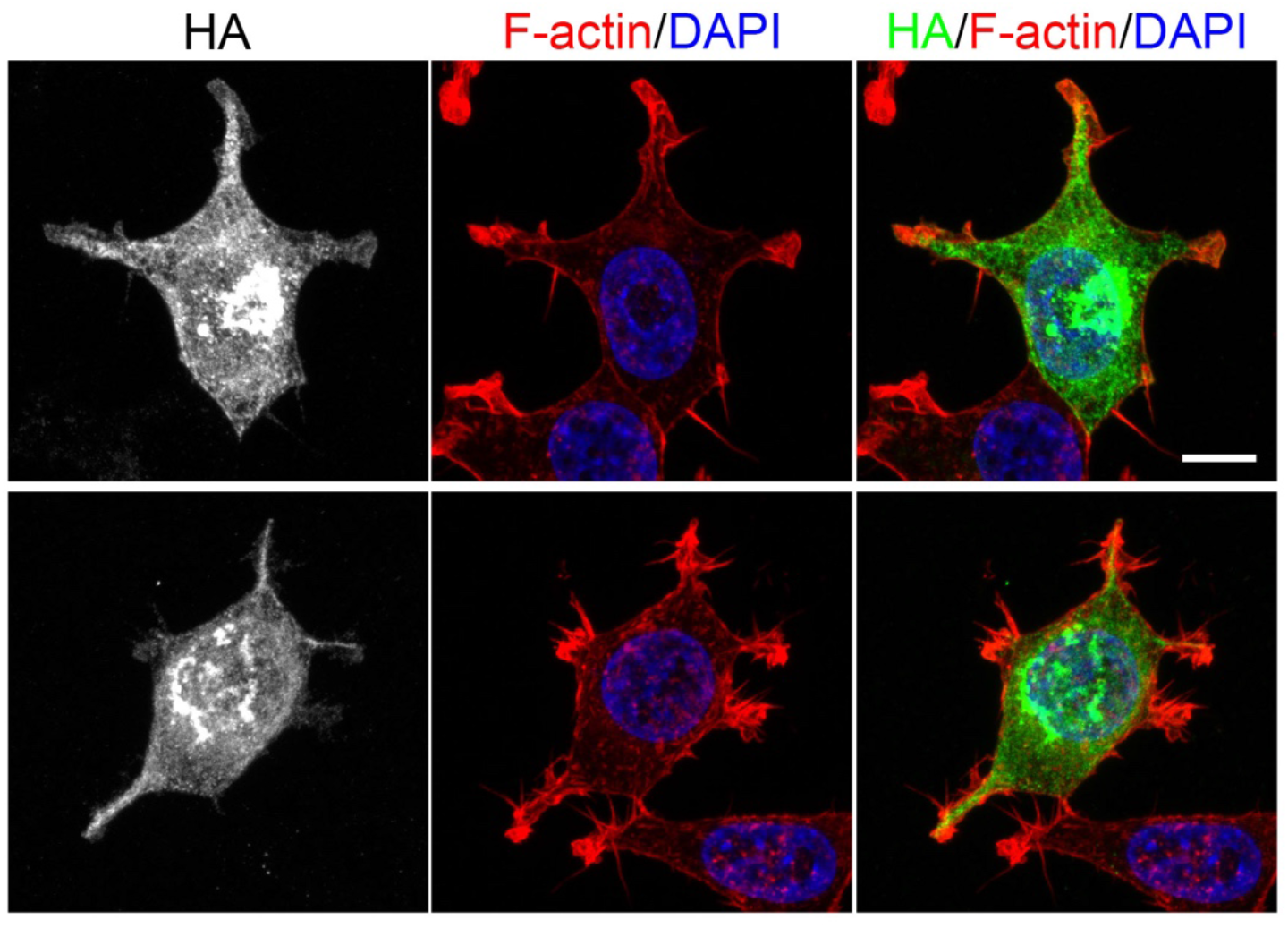
TLR3 is not enriched in F-actin-positive structures. C-terminal HA-tagged TLR3 was expressed in HEK293T cells. Cells were immunofluorescently stained with HA antibody. F-actin was labeled using Alexa-fluor-546-conjugated phalloidin. The nuclei were counterstained with DAPI. Scale bars: 10 μm.

